# Global biogeographical regions reveal a signal of past human impacts

**DOI:** 10.1101/586313

**Authors:** Marta Rueda, Manuela González-Suárez, Eloy Revilla

## Abstract

Ecologists have long documented that the world’s biota is spatially organized in regions with boundaries shaped by processes acting on geological and evolutionary timescales. Although growing evidence suggests that human impact has been key in how biodiversity is currently assembled, its role as a driver of the geographical organization of biodiversity remains unclear. Here, we quantify the relative importance of human land use from ∼5000 years ago to predict the current assemblage of terrestrial mammals in biogeographical regions across the Earth. Results show that past anthropogenic land use has left an imprint on the taxonomic differentiation of some of the largest biogeographical realms, whereas land use at present stands out as a driver of the taxonomic differences between medium-sized subregions, i.e., within and among continents. Our findings highlight the far-reaching effect that past anthropogenic actions have had on the organization of biodiversity globally.

## Introduction

How is the world’s biodiversity organized, and why do large-scale patterns of taxonomic diversity change through natural geographic regions? These questions have attracted the attention of naturalists since the early 19^th^ century (von Humboldt, 1806; Wallace, 1876; Ricklefs, 2004; Daru et al., 2017) and are now critical to forecast the future of biodiversity in the face of global change (Botkin et al., 2017). A key step in understanding the organization of biodiversity is to identify the assemblage of regions based on their shared organisms (Wiens, 2011). Alfred R. Wallace was among the first to propose that the world’s fauna is organized hierarchically in broad regions shaped by geographic and climatic factors (Wallace, 1876). About 150 years later, the development of multivariate analytical techniques has led to revaluations of Wallace’s proposal (Kreft and Jetz, 2010; Procheş and Ramdhani, 2012; Holt et al., 2013; Rueda et al., 2013) and the refinement of our understanding regarding the extrinsic determinants driving the main dissimilarities among bioregions (Ficetola et al., 2017). Broadly, what we know is that the interplay of multiple factors has jointly contributed to the formation of bioregions, but their prominence varies across the globe (Riddle and Hafner 2010; Ficetola et al., 2017; Mazel et al., 2017). Processes acting deeper in the past, like plate tectonics, are the most important in explaining the separation between strongly divergent biogeographical realms. Factors representing present-day ecological barriers, like climatic heterogeneity, determine the separation between less dissimilar regions, i.e., within and among continents; while mountains - the third factor in importance - operate at all levels of biogeographical differentiation (Ficetola et al., 2017). However, although there is clear evidence that anthropogenic actions are the main driver of the extinction processes acting at multiple scales (Barnosky et al., 2011; Ceballos et al., 2017) and, thereby, of the biotic homogenization of the ecological communities worldwide (Olden, 2006; Baiser et al., 2012), we do not yet fully understand whether and how anthropogenic impact has caused changes at a biogeographical level.

Anthropogenic impact has usually not been contemplated in the biogeographical scenario likely because extant biogeographical regions have traditionally been assumed to reflect the natural organization of biodiversity that resulted from ecological, historical, and evolutionary processes acting over millions of years (Lomolino et al., 2010). However, rising evidence showing how human-mediated species introductions are already affecting extant biogeographic regions (Capinha et al., 2015; Bernardo-Madrid et al., 2019) challenges this view. Furthermore, accumulating facts indicate that Quaternary human activities induced shifts in the plant and animal communities we see today (Lyons et al., 2016), and there is little doubt that modern humans have been a major driver in the extinction of large mammals during the late Pleistocene and early Holocene (Sandom et al., 2014; Smith et al., 2018). Moreover, archaeological, palaeoecological and genetic data suggest that cumulative human transformation of ecosystems over millennia has resulted in dramatic changes in composition, community structure, richness and genetic diversity of a diverse array of organisms across taxonomic groups (Ellis et al., 2013, 2021; Boivin, 2016; Mottl et al., 2021). This body of evidence raises the question of whether historical anthropogenic pressures may have been large enough to leave an imprint detectable today on biogeographical assemblages globally.

To answer these questions, we set up a bioregionalization and analytical procedure focused on two data sets; one describing the current distribution ranges of global terrestrial mammals, and another set inferring their present natural ranges, which represents estimates of where species would potentially live without anthropogenic pressures (Faurby and Svenning, 2015; Faurby et al., 2018). Using both types of distribution ranges we build hierarchical bioregionalizations, upscaling from the smallest detectable bioregions to the largest realms, and compute models to identify those variables (plate tectonics, climatic heterogeneity, mountains and anthropogenic land use over the Holocene, specifically 5000 and 2000 years ago as well as present time) that best predict the assemblage of species in bioregions. We hypothesized that if anthropogenic land use has had a noteworthy effect on the biogeographical assemblages and boundaries as we know them nowadays, then (1) the biogeographic configurations built using current distribution ranges will differ from those based on natural ranges; and (2) human land use should predict the biogeographic patterns obtained using current ranges, but not patterns obtained using natural ranges. To reinforce our results, we also run complementary models using only the distribution ranges of those extant terrestrial mammals that have undergone some modification in their range due to anthropogenic pressures (536 species), i.e. they have an estimated distribution range in PHYLACINE different from that of the IUCN (2016) (Faurby et al., 2018). Besides, at the largest biogeographical scale we also compute models excluding Madagascar and Australia as these bioregions are characterized by an extremely low historical land use, which could influence results.

Our analyses reveal that historic anthropogenic land use have left an imprint on how diversity is structured on Earth at a biogeographical scale and are consistent with increasingly evidence showing that the impact of past human actions on ecosystems and organisms has been greater than previously thought.

## Results

### Bioregionalizations

Both for the current and natural bioregionalizations we identified a hierarchical system of biogeographic regions with four levels (Appendix 1-Figure 1). For current mammalian ranges, the broadest delineation of nine large bioregions was mostly consistent with the original maps of Wallace’s realms and subregions (Figure 1a). For natural mammalian ranges, the eight bioregions obtained showed a different biogeographic arrangement with respect to the nine large current ones, especially in the Northern Hemisphere (Figure 1b) (Table 1). Differences between current and natural bioregions (‘Material and Methods’) extended to the subregions or medium-sized bioregions and smaller bioregions (Appendix 1-Figure 1) and were confirmed for the bioregionalizations carried out with the 536 species whose distribution ranges have been modified by anthropogenic actions (Appendix 1-Figure 2 and Appendix 1-Table 1).

**Fig. 1.**
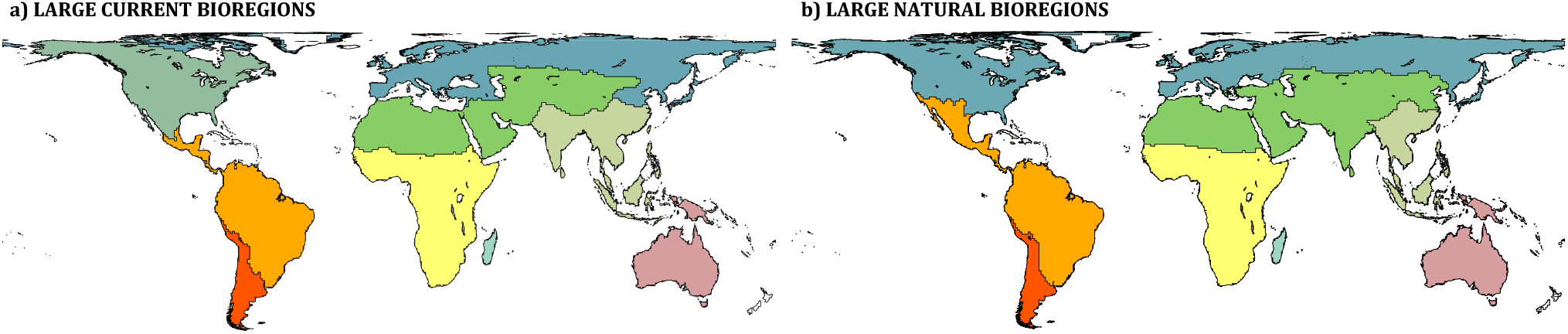
Current and natural broader regions or realms. Only results for the broader regions are shown here: 9 current bioregions (a), and 8 natural bioregions (b). The greatest differences between both biogeographical configurations are more relevant in the Northern Hemisphere, where for the natural bioregions we found that the Palearctic and the Nearctic are united forming a great extension, the typical biographical boundary between the Nearctic and the Neotropical realms disappears and extends towards the north, and India is no longer part of the Asian region and is included instead within the ‘Saharo-Arabian’ region.

**Table 1.** Values of Jaccard index between current and natural hierarchical bioregions. Jaccard index assesses the degree of similarity between clusters (bioregions) and ranges between 0 (no similarity) and 1 (perfect match). The number of bioregions obtained during each bioregionalization processes is also provided.

| CURRENT | Jaccard<br>coefficient | NATURAL |
| --- | --- | --- |
| 9 broader realms | 0.57 | 8 broader realms |
| 27 subregions | 0.58 | 25 subregions |
| 141 small bioregions | 0.42 | 121 small bioregions |
| 1128 smaller bioregions | 0.28 | 1051 smaller bioregions |

**Fig. 2.**
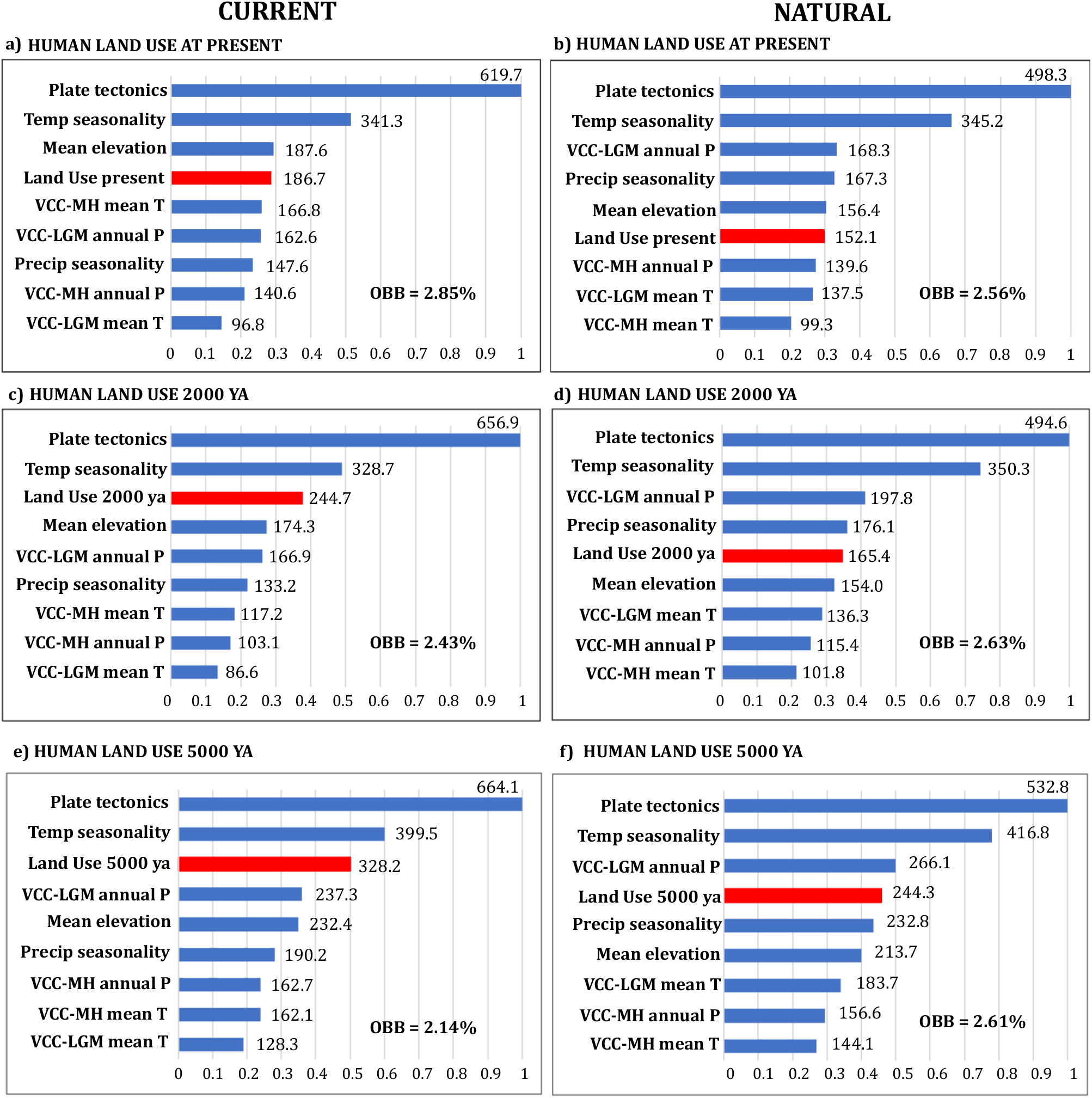
Ranking of importance values for the drivers of taxonomic differentiation for the current and natural broader bioregions. Above panels show importance values from models with human land use at present for (a) current and (b) natural bioregions, while the panels below show importance values from models with human land use 2000 years ago for (c) current and (d) natural bioregions, and human land use 5000 years ago for (e) current and (f) natural bioregions. Importance was measured by the drop-in classification accuracy after predictor randomization in random forests of 5000 trees. Higher values of mean decreased in accuracy indicate variables that are more important to the classification. OBB (Out-of-bag) represents the percentage of cells misclassified in each model. VCC = velocity of climate change; MH = Mid-Holocene; LGM = Last Glacial Maximum; annual P = annual precipitation; mean T = mean annual temperature; ya = years ago.

### Global importance of predictors of the biogeographical patterns

For the broader bioregions, past human land use (5000 and 2000 years ago) was the third most important predictor of the biogeographic structure of current realms behind plate tectonics and seasonality in temperature and ahead of mountains (Figure 2c, 2e), with a moderate but non-negligible predictive value (∼22% and 21% of cells classified correctly when the model was run just with human land use 2000 and 5000 years ago, respectively, see Figure 3). Human land use at present had less relevance in the classification models (Figure 2a). In accordance with our prediction, past and present human land use were poorer predictors of natural bioregions (Figure 2b, d, f). Results were consistent also if we excluded Australia and Madagascar, two biogeographic regions with minimal human land use before 2000 years ago (Appendix 1-Figure 3). Models run with the unique 536 extant terrestrial mammals whose range of distribution has been affected by human impact revealed present and past human land use as the second predictor but with negligible importance with respect to plate tectonics, which was by far the most important driver (Appendix 1-Figure 4). The misclassification rate (Out-of-bag, OBB hereafter) for these set of models was quite low, never exceeding 3%.

**Fig. 3.**
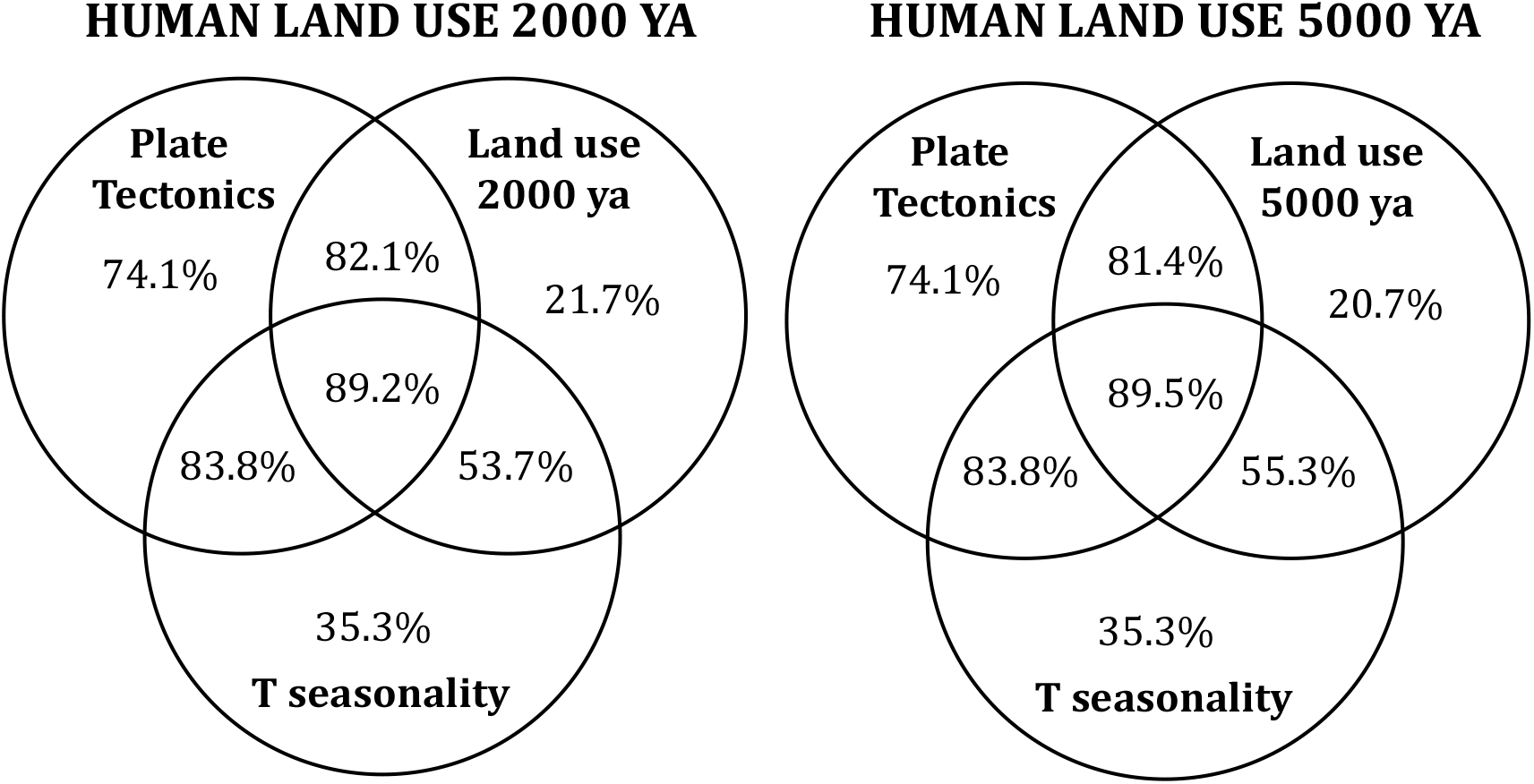
Diagrams describing the accuracy in classification of the three most important drivers of the current broader bioregions for models including human land use 2000 and 5000 years ago. Values indicate the percentage of cells correctly classified (1-OBB error) in their bioregion, and are obtained from running random forest classification models for each individual variable, by pairs and for the three most important factors together.

**Fig. 4.**
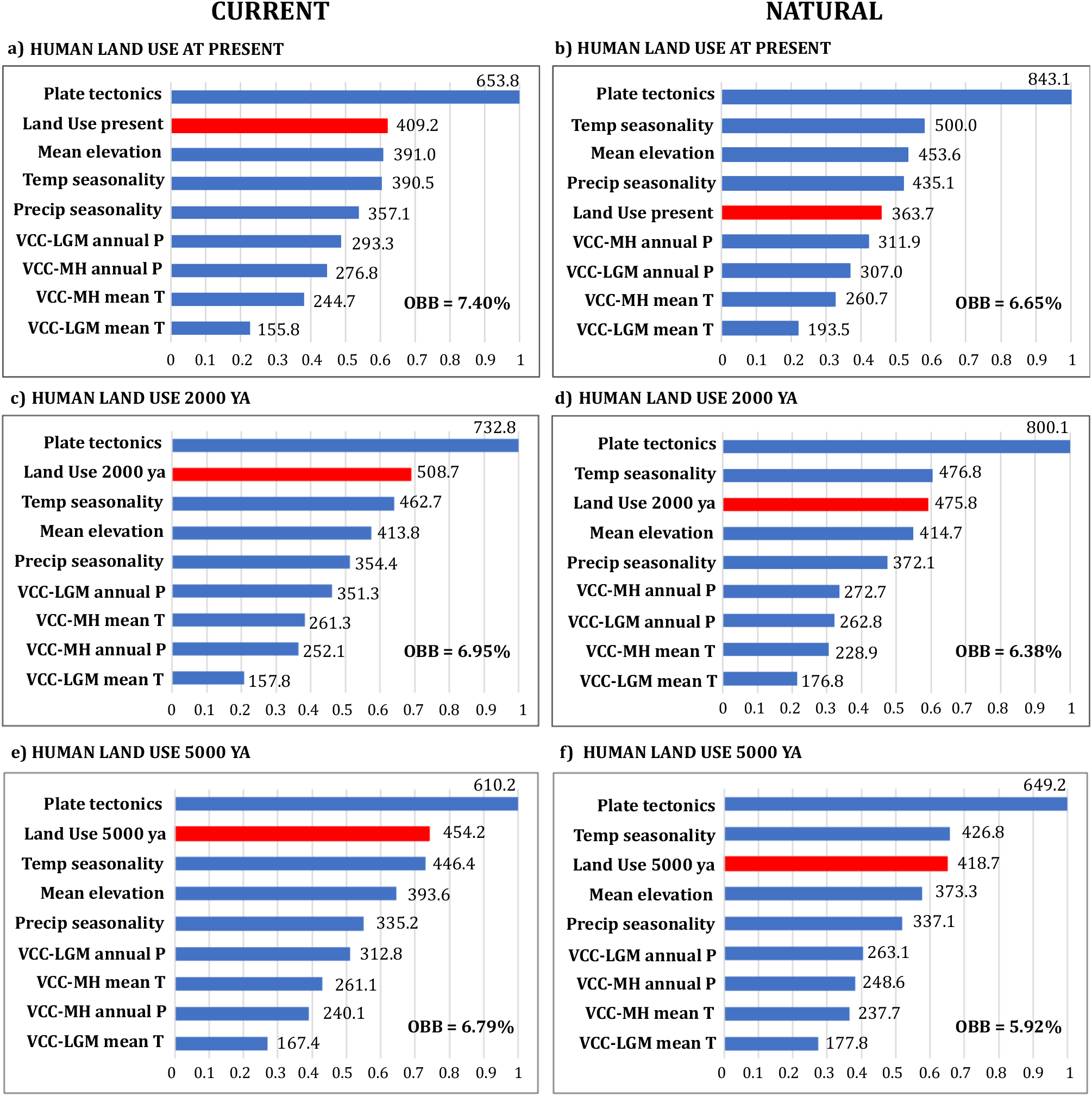
Ranking of importance values for the drivers of taxonomic differentiation for the current and natural medium-sized bioregions or subregions. Above panels show importance values from models with human land use at present for (a) current and (b) natural bioregions, while the panels below show importance values from models with human land use 2000 years ago for (c) current and (d) natural bioregions, and human land use 5000 years ago for (e) current and (f) natural bioregions. Specifications as in Fig. 2.

For medium-sized bioregions or subregions, past and present human land use were the second predictors in importance for the current bioregions (Figure 4 a, c, d; OBB values in general < 7.50%). As hypothesized, the importance of land use was smaller for the natural bioregions, although its past land use remained a relatively strong driver (Figure 4 b, d, e). Results were consistent for the models run with the unique 536 extant terrestrial mammals influenced by human actions (Appendix 1-Figure 5; OBB < 8.5%).

**Fig. 5.**
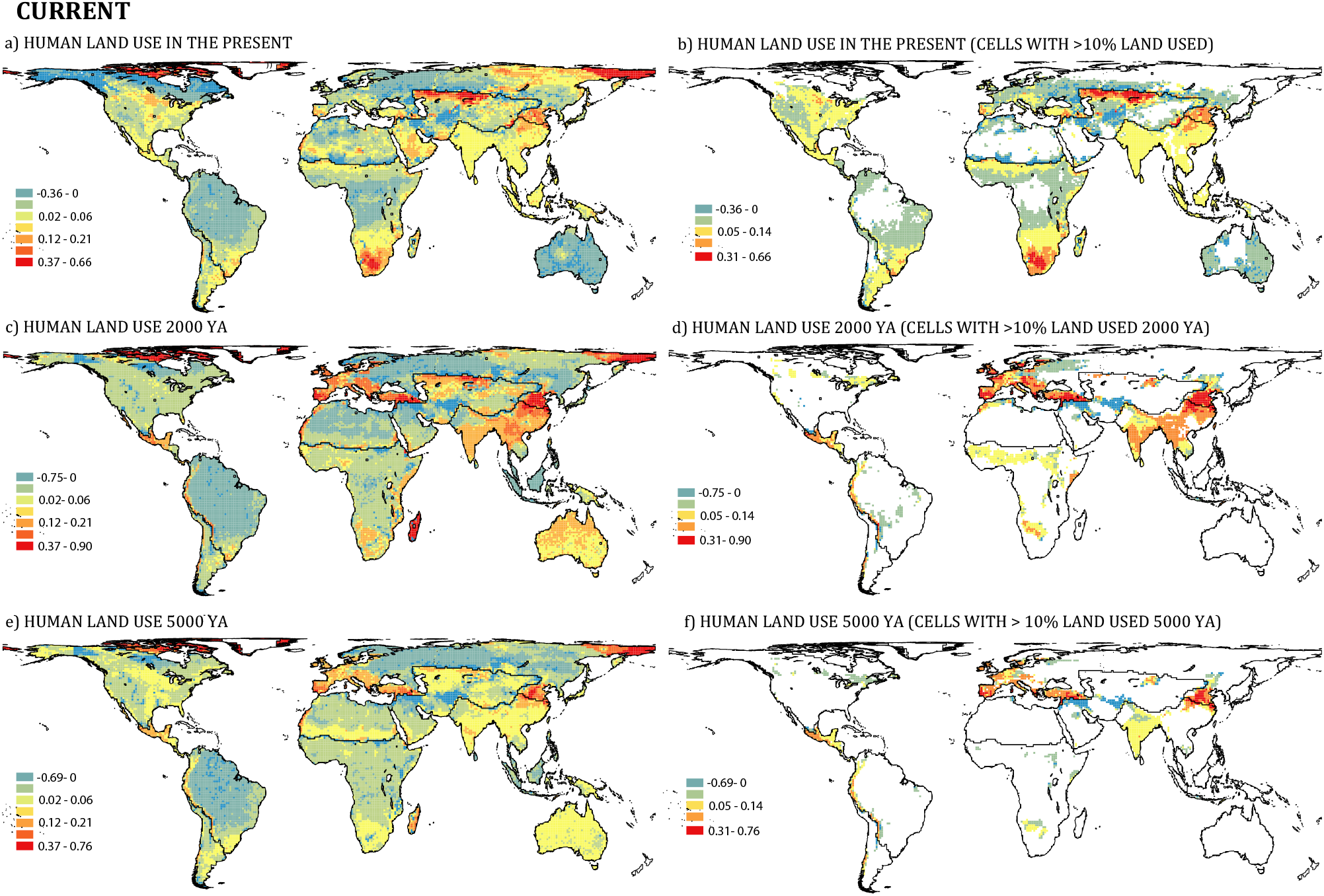
Maps of the local importance of present and past human land use for the current broader bioregions. The left panels represent the local importance values for human land use in the present (a), 2000 years ago (c) and 5000 years ago (e). The right panels represent local importance values when land use at present (b), 2000 (d), and 5000 years ago (f) was >10% in each grid cell. The local importance shows the importance of the variable (human land use here) for the classification of a single sample (grid cells). The score of the legends shows the impact on correct classification of grid cells, with blue colors indicating from negative to 0 values (incorrect to neutral classification), and yellow, orange and red indicating positive values (correct classification). Since the values of local importance do not distinguish cells where the human land use was significant and areas where it was negligible, i.e., both can be important for the classification process (‘Material and Methods’), we have also mapped the importance values for those grid cells where the percentage of land use at present, as well as 2000 and 5000 years ago was >10% for a better interpretation. In these maps, white cells imply human land use values <10%.

As bioregions became smaller, many more variables were necessary to predict the classification of cells into bioregions, and results for the current and natural bioregions became quite similar (Appendix 1-Figure 6 and 7; OBB < 15.1% and < 35.6% for small and very small bioregions, respectively). Overall, human land use at present has minor relevance in the classification models, whereas past human land use remained relatively high, being third or fourth in importance (Appendix 1-Figure 6 and 7).

**Fig. 6.**
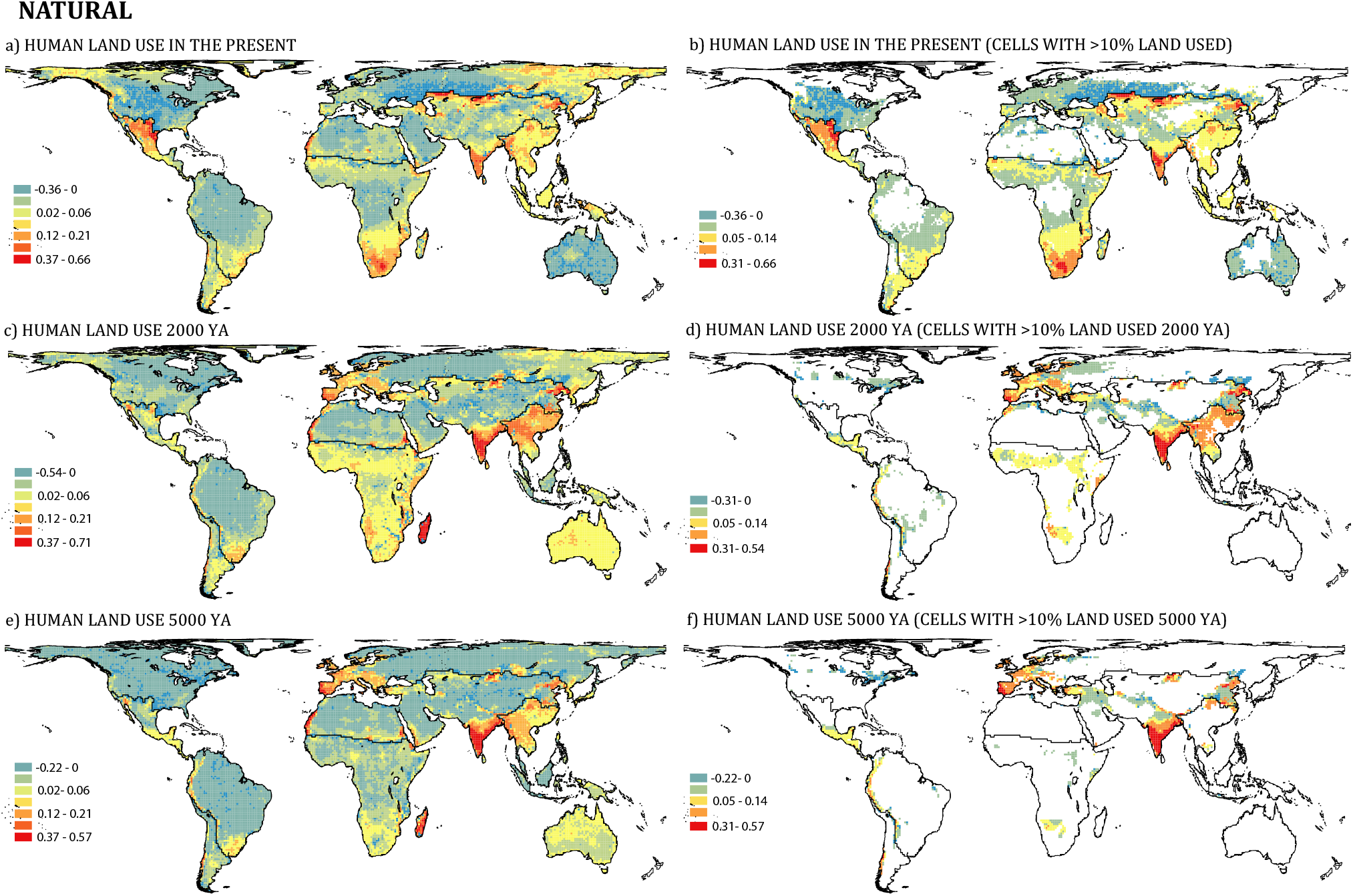
Maps of the local importance of present and past human land use for the natural broader bioregions. The left panels represent the local importance values for human land use in the present (a), 2000 years ago (c) and 5000 years ago (e). The right panels represent local importance values when land use at present (b), 2000 (d), and 5000 years ago (f) was >10% in each grid cell. The score of the legends shows the impact on correct classification of grid cells, with blue colors indicating from negative to 0 values (incorrect to neutral classification), and yellow, orange and red indicating positive values (correct classification).

**Figure 7.**
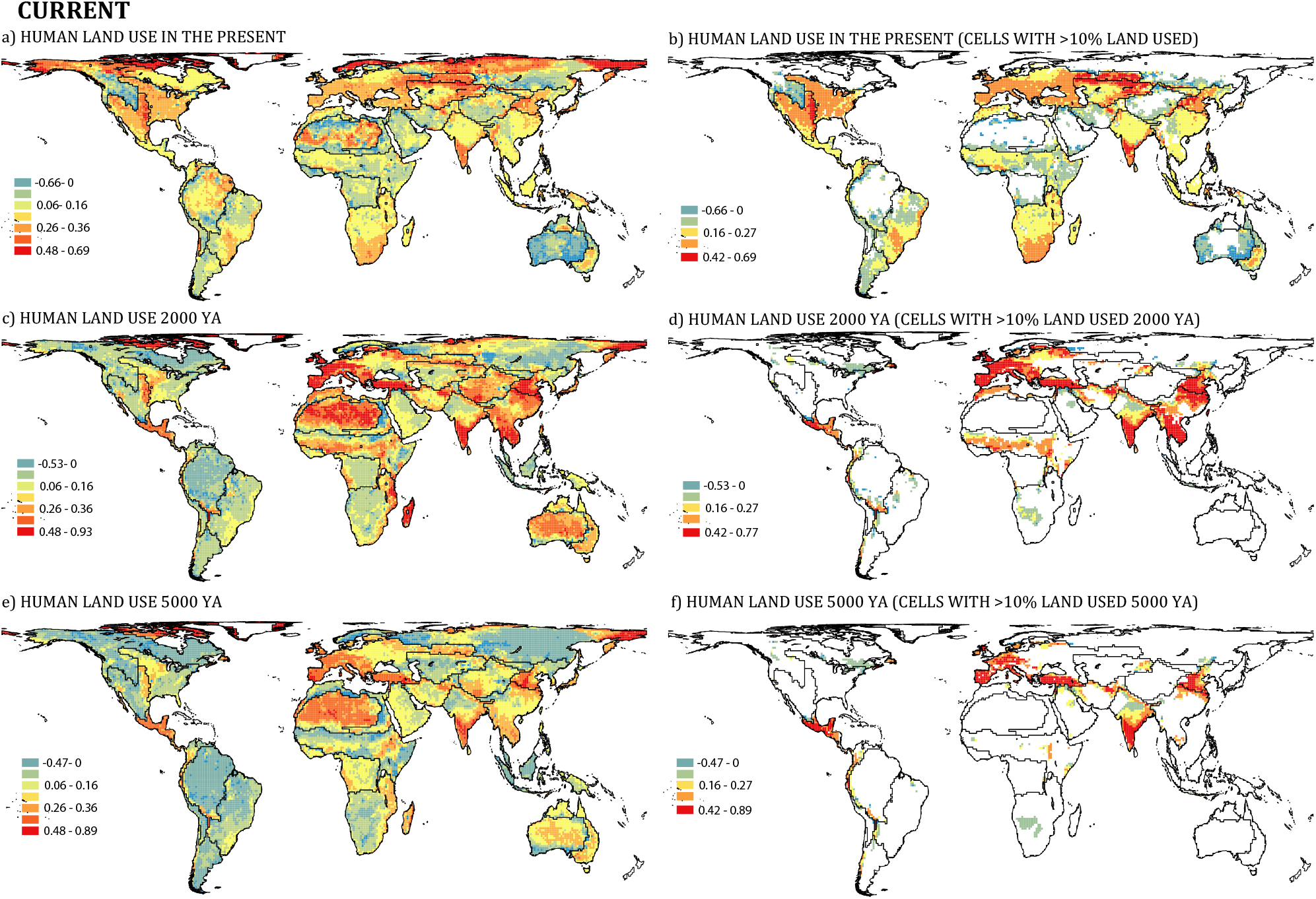
Maps of the local importance of present and past human land use for the medium current bioregions. The left panels represent the local importance values for human land use in the present (a), 2000 years ago (c) and 5000 years ago (e). The right panels represent local importance values when land use at present (b), 2000 (d), and 5000 years ago (f) was >10% in each grid cell. The score of the legends shows the impact on correct classification of grid cells, with blue colors indicating from negative to 0 values (incorrect to neutral classification), and yellow, orange and red indicating positive values (correct classification).

### Local importance of predictors of the biogeographical patterns

Maps showing local importance scores (‘Material and Methods’) revealed large differences in the location of grid cells where past and present human land use have been analytically important for the classification of the broader current bioregions (Figure 5). The cells where human land use at present was most relevant are located in southern Africa, the easternmost boundary between the Palearctic and the Asian regions, part of the northern boundary between the Palearctic and ‘Saharo-Arabian’ regions, and much of the arctic zone of the Palearctic (Figure 5a), although note that grid cells of the two latter areas are important because of their lack of human land use (Figure 5b) (‘Material and Methods’). The cells where human land use 2000 years ago was more important are located mainly throughout Central America and the Andes mountain range, Europe except for the northernmost part, most of the Asian region, the boundary between the Palearctic and the Asian regions (Figure 5c), as well as Madagascar and much of the arctic zone of the Palearctic but due to lack of human land use (Figure 5d). The cells where human land use 5000 years ago was more important mostly coincided with those of 2000 years ago, but with minor values of importance (Figure 5 e, f). All these areas mostly coincided for the broader natural bioregions (Figure 6).

For the medium current bioregions, the grid cells where the human land use at present was most relevant for the classification of bioregions included those highlighted for the broader bioregions plus most part of Europe, southern India, eastern Australia, a large area of the Nearctic, and the Sahara Desert (Figure 7 a, b). When focusing on the human land use 2000 years ago, the areas affected by human land use were similar to those of the broader bioregions, but note that other areas also became very important (e.g. Sahara Desert, Mongolian Plateau) due to their lack of human land use (Figure 7 c, d). Again, the cells where human land use 5000 years ago was more important mostly coincided with those of 2000 years ago but with minor importance values (Figure 7 e, f). The location of cells affected by human land use for the medium current bioregions quite coincided with that for natural bioregions (Figure 8).

**Figure 8.**
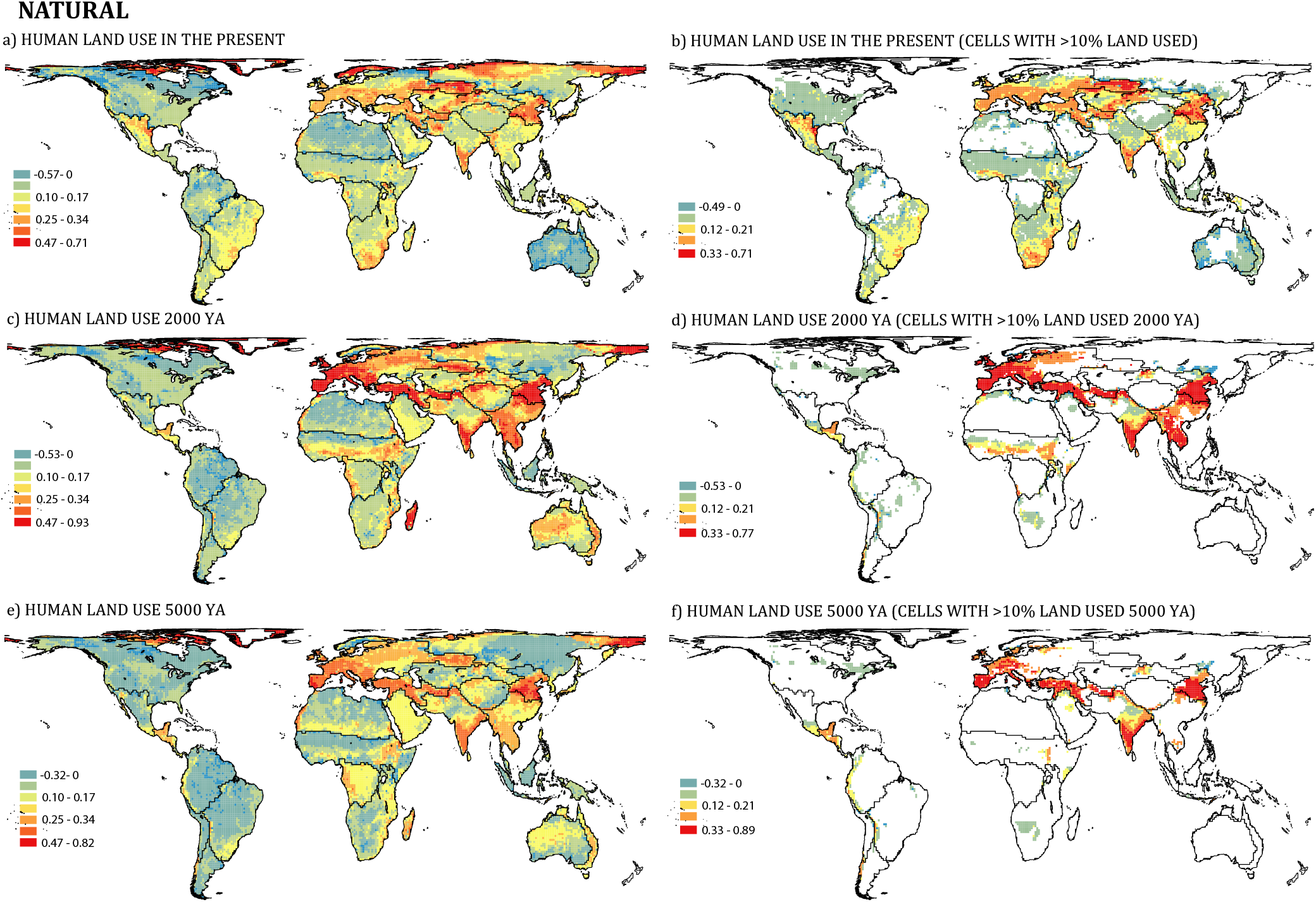
Maps of the local importance of present and past human land use for the medium natural bioregions. The left panels represent the local importance values for human land use in the present (a), 2000 years ago (c) and 5000 years ago (e). The right panels represent local importance values when land use at present (b), 2000 (d), and 5000 years ago (f) was >10% in each grid cell. The score of the legends shows the impact on correct classification of grid cells, with blue colors indicating from negative to 0 values (incorrect to neutral classification), and yellow, orange and red indicating positive values (correct classification).

## Discussion

Our study shows that human activity during the last few millennia has left its mark on the spatial organization of the Earth’s biodiversity. First, we show that biogeographical configurations based on current and natural mammal distributions differ, particularly in the Northern Hemisphere. These differences were not solely the result of including extinct megafauna - with huge distributions ranges – in the natural biogeographic regions, as biogeographical patterns for the 536 extant terrestrial mammals whose ranges of distribution have been affected by anthropogenic impacts confirmed our findings. Second, we show that human land use is an important predictor of current bioregions but does not largely contribute to predict natural bioregions. For example, for the broader regions or realms, past human land use was the third most important predictor behind plate tectonics and temperature seasonality and ahead of mountains, but has minor importance on natural regions. Past anthropogenic actions have seemingly left a perceptible biogeographical imprint on the assembly of the global realms we recognize today. Moreover, for the medium-sized subregions, human land use at present was the second most important predictor suggesting that recent human activity is already causing changes in the biogeographical assemblages at the subregional scale, i.e., within and among continents.

The mammalian distributions that we know today are not only a reflection of the most recent human actions but also those exerted during the last few millennia (Faurby and Svenning, 2015; Polaina et al., 2020). Recent studies had already shown that human actions in the present, in terms of human-mediated species introductions, can be evident at biogeographical scales (Capinha et al., 2015; Bernardo-Madrid et al., 2019), but here we also show evidence of an effect of past human land use. What did happen 5000 years ago that left such as noticeable footprint in the Earth’s biogeographical regions? This period coincides with the emergence of an “intense” agricultural land expansion and the settlement of densely populated areas in the Mediterranean, Southwest Asia, South Asia, and eastern China (Kirck, 2005; Ellis et al., 2013; Stephens et al. 2019). Long-term impacts to enhance agricultural productivity, including forest clearing, increased fire frequencies, species invasions, and soil erosion, were already apparent in some regions at this time (Ellis et al., 2013; Boivin et al., 2016), and may have transformed vegetation structure and species composition across many regions, from the Mediterranean to the Tropics (Stephens et al. 2019). In fact, reconstructions suggest that >20% of Earth’s temperate woodlands had been already impacted by humans by the late-Holocene (Ellis et al., 2013). Indeed, our results on local importance show the match between some of the biogeographical boundaries and regions of the New World and the distribution – and presumed impact – of ancient Mesoamerican and South American pre-European civilizations (Richard, 1997; Dunning et al., 2013). In the Old World, we note the prominence of past human land use covering most of the Asian realm territory, likely associated to the rise of rice cultivation in eastern, southern and southeast Asia over the Holocene (Silva et al., 2015). Indeed, cumulative archaeological data show that the scale of agriculture and land use in these regions was significant; expansion of land area used for livestock and rice (*Oryza sativa*) paddy agriculture in southern Asia was sufficient to increase atmospheric methane emissions between 4000 and 1000 years ago (Fuller et al., 2015). Moreover, the atmospheric CO_2_ decline registered between A.D. 1570 and 1620 has been attributed to the uptake by vegetation following the agriculture abandonment caused by the population crash after the arrival of Europeans to America (Lewis and Maslin, 2015). Past human land use can also partly explain differences between the southern Palearctic boundary and the Afrotropic below the Sahara. Both can be related to the transformation of European and Near Eastern landscapes during the Roman period (Butzer, 2005), and the expansion of sorghum (*Sorghum bicolor*) cultivation 3000 to 2000 years ago in the sub-Saharan Africa (Boivin et al., 2016).

While we found differences in importance of human land use 5000 and 2000 years ago between current and natural medium-sized bioregions, we were surprised that past human land use was an important determinant of natural bioregions, which were based on distributions of mammals in the absence of human impact. These natural ranges are inferred ‘natural’ distributions and it is possible that they did not fully exclude human impacts, as for 3424 of the extant mammals analyzed, natural and current distributions were identical. In addition, these results could also reflect an overlap between human land use and biogeography. If the distribution of humans during the middle and late Holocene responded to the same drivers as the distributions of terrestrial mammals, human land use could appear as a predictor even if it had no causal effect. In fact, the importance of this variable appears to be a consequence of how human land use was distributed globally 5000 and 2000 years ago, or more accurately where human activity was largely absent. Note that while current human land use is distributed over wide areas of the planet, that of 5000 and 2000 years ago was centralized in specific areas coinciding with the distribution of ancient civilizations. This means that on a subregional scale, many bioregions can be well predicted by the absence of human land use. Indeed, an additional analysis that excluded subregions with no reported human land use 2000 years ago showed very little importance of past human land use for both current and natural medium-sized bioregions (Appendix 1-Figure 8).

In summary, our results show the value of considering impacts of past human actions to understand the current organization of biodiversity globally. Previous studies have documented lasting effects of human land use changes during the last millennia on current biodiversity patterns (Dambrine et al, 2007; Haberle, 2007; van der Sande et al., 2019), but this is the first time the signal has been recognized on the taxonomic differentiation of the largest realms. This would mean that some of the biogeographical boundaries proposed by Wallace 140 years ago (Wallace, 1876) were already showing the influence of anthropogenic actions. The human transformation of ecosystems that occurred in large terrestrial extensions coinciding with the distribution of ancient civilizations, makes past human land use important along with orographic processes and temperature seasonality to discern taxonomic differences among the large biogeographical zones of the world. It is certainly difficult to disentangle direct causal effects from correlative relationships in non-experimental large-scale studies such as this. We know the same geographic and environmental characteristics that influenced the distribution across space and time of mammals likely also influenced the distribution of our ancestors (Lomolino, 2018). However, we considered several of those predictors and still found a global imprint of past human actions on the patterns of distribution of global diversity. Our results force us to reflect on how the much more widespread and severe changes that have occurred since the beginning of the industrial revolution will affect the organization of biodiversity in the future. We show here the effects of recent human actions are already detectable at the subregional scale, and possible at the broader biogeographical scale. Although it may seem like an unlikely or distant future, the possibility that human activities become the main driver, above historical geomorphological or climatic processes, that shapes Earth’s biodiversity cannot be ignored.

## Material and Methods

### Generating presence-absence matrices using range distributions

Current and natural distribution ranges of terrestrial mammals were obtained from PHYLACINE 1.2 (Faurby et al., 2018), which contains range maps for all 5831 known mammal species that lived since the last interglacial (∼130,000 years ago until present). Ranges for current extant species contained in PHYLACINE are IUCN (2016) distribution maps, whereas present natural ranges represent estimates of where species would potentially live if they had never experience anthropogenic pressures, and – in the case of extinct species – had not gone extinct (see Faurby and Svenning, 2015 for details). Both current and natural ranges in PHYLACINE are projected to Behrmann cylindrical equal area rasters with a cell size of 96.5 km by 96.5 km at 30° North and 30° South. We generated two presence-absences matrices, one for the current and one for the natural ranges, in which every row represents a grid cell and every column a species. From theses matrices we excluded (1) *Homo* species; (2) bats and marine mammals; (3) cells containing less than 50% of land area to approximate equal-size samples, and (4) cells containing fewer than five species to reduce potential distortions caused by having few taxa (Kreft and Jetz, 2010; Rueda et al., 2013). These exclusion criteria rendered a total of 14,087 cells representing 3960 terrestrial mammal species for the current matrix, and 14,151 cells representing 4306 for the natural matrix. With the aim of conducting complementary analyses to support our results we also generated two additional matrices using only those extant mammals that have undergone some modification in their distribution ranges due to anthropogenic pressure, i.e., they have an estimated distribution range in PHYLACINE different from that of the IUCN, representing 536 species.

### Building biogeographical regionalizations

To each of the presence-absence matrices we applied a machine-learning algorithm referred to as affinity propagation (AP hereafter) (Frey and Dueck, 2007) to build biogeographical regionalizations at different biogeographical resolution. We chose AP because it can compress massive data sets very efficiently (i.e. with lower error), and its good performance in hierarchical bioregionalization procedures (from few grid cells to large realms) has already been demonstrated in a previous work (Rueda et al., 2013). The AP algorithm works detecting special data points called exemplars, and by a message-passing procedure it iteratively connects every data point to the exemplar that best represents it until an optimal set of exemplars and clusters emerges. Contrary to algorithms in which exemplars are found by randomly choosing an initial subset of data points, AP takes as input measures of ‘similarities’ between pairs of data points (grid cells here) and simultaneously considers all the points as potential exemplars. The optimal set of exemplars is the one for which the sum of similarities of each point to its exemplar is maximized. Hence, detecting exemplars goes beyond simple clustering because the exemplars themselves store compressed information (Mézard, 2007).

We first used the presence-absence matrices to calculate pairwise similarities between pairs of cells, and selected Hellinger distance as a similarity index. The Hellinger distance is a modification of the Euclidean distance (Legendre and Gallagher, 2001) and is used to avoid the ‘double-zero’ problem, i.e. when two sites that have no species in common are assigned the same distance as two sites that share the same species; and the ‘species-abundance paradox’, which frequently occurs when two sites share only a small fraction of all the species in the same regional pool. This is expected to be a particular problem at the margins of biogeographical regions where sites may be quite different from one another rather than in the centre of a region where sites are likely to be very similar in their species assemblages (Legendre and Legendre, 1998; Gagné and Proulx, 2009). We then used a protocol based on a successive application of AP (Rueda et al., 2013) to obtain a biogeographical upscaling from the smallest possible bioregions (i.e. the highest biogeographical resolution) to the largest ones. We performed an initial AP analysis involving all grid cells of the similarity matrix. This first AP run generates the optimal solution of the highest resolution bioregions, while also identifying its exemplars. We obtained 1128 clusters and exemplars for the current distributions and 1053 for the natural distributions. Then, using the exemplars as the new units of analysis we conducted again an AP, i.e., we calculated a new similarity matrix and re-run a new AP. This process was repeated until a small and coherent number of large clusters emerged. Finally, to obtain maps of each clustering result, we classified each grid cell (row) of every presence-absence matrix according to the cluster to which they were assigned in its corresponding AP analysis. AP analyses were performed using the ‘*APCluster*’ R package (Bodenhofer et al., 2001).

### Assessing differences in biogeographical configurations

The degree of similarity between clusters (i.e. regions) of current and natural bioregions was estimated using the Jaccard index. For that, we used the *cluster_similarity* function of the ‘*clusteval*’ R package (Ramey, 2012). This function computes the similarity between two clusterings of the same data. Jaccard index ranges between 0 (no similarity) to 1 (perfect match).

### Predictors

To each of the cells of the presence-absence matrices we assigned a mean value of several predictors. We considered variables previously tested as determinants of biogeographical boundaries related to climatic heterogeneity, orographic barriers, tectonic movements, and instability of past climate (e.g. Ficetola et al., 2017) plus variables related to anthropogenic land use over the Holocene. In particular, we tested four climatic variables: annual total precipitation, mean annual temperature, seasonality in temperature and seasonality in precipitation. All climatic variables were extracted from the WorldClim dataset (Fick and Hijmans, 2017) up-scaled at a 96.5 km resolution. These variables represent both average conditions and variability within years and have been shown to be determinants of vertebrate distributions (Sexton et al., 2009). Climatic conditions have strongly shifted over the Quaternary period, and have been shown to play a role in the present-day species distributions, endemism, and assemblages (Graham, 1996; Davis and Shaw, 2001; Araujo et al., 2008; Sandel et al., 2011). To test for the potential effect of past climate change or stability, we calculated the average velocity of climate change since the Last Glacial Maximum (LGM; ∼22,000 years ago) and since the Mid Holocene (MH; ∼ 6000 years ago) (see Sandel et al., 2011). For that, we used the mean annual temperature and annual total precipitation for the MH and the LGM calculated by means of the Model for Interdisciplinary Research on Climate (MIROC-ESM) (Watanabe et al., 2011). Mountain ranges represent major barriers to dispersal for most mammals, whereas plate tectonics are responsible of the long-term isolation of the biotas (Ficetola et al., 2017, Lomolino et al., 2010). We used the GTOPO30 to calculate the mean elevation per grid cell. Plate tectonics were obtained from Bird (2003). Each grid cell was assigned the tectonic plate to which it belongs. When a cell was represented by more than one tectonic plate, it was assigned the one that occupied a greater percentage of the cell. Finally, past anthropogenic land use was obtained from Ellis et al. (2013). This dataset consists of two spatially explicit global reconstructions of land-use from two main models of land use across the Holocene, HYDE and KK_10_ models. Among them, we chose the more realistic KK_10_ model, which assumes that humans use land more intensively when population density is high and land scarce (Kaplan et al., 2011). In counterpoint, the HYDE model omits land-use intensification and predicts that except for the developed regions of Europe, human use of land was insignificant in every biome and region before A.D. 1750. This dataset comprises raster information for human land use for 10 different time periods; B.C. 6000, 3000, and 1000, and A.D. 0, 1000, 1500, 1750, 1900, 1950 and 2000. We calculated human land use at four different time spans: 8000, 5000, and 2000 years ago, as well as at the present time (A.D. 2000). However, we decided do not include land use 8000 years ago in the random forest models due to its low representation at global level (Appendix 1-Figure 9).

### Evaluating potential drivers of the biogeographical configurations

We used random forest (Breiman, 2001) models of 5000 classification trees to assess the factors that may predict the classification of cells in bioregions at different biogeographical resolutions and to estimate the relative importance of the predictors. Random forest is a machine learning method based on a combination of a large set of decision trees. Each tree is trained by selecting a random set of variables and a random sample from the training dataset (i.e., the calibration data set). The accuracy of the models is given by the out-of-bag (OOB), an estimate of the misclassification rate that represents and unbiased error rate of the model that is calculated by counting how many cases in the training set are misclassified and dividing the number by the total number of observations.

Random forests are able to disentangle interacting effects and identify nonlinear and scale-dependent relationships that often occur at the scale of the analysis performed here among multiple correlated predictors (Cutler et al., 2007). Although random forests are generally assumed to not be affected by highly correlated predictor variables, we eliminated some predictors showing a moderate to high correlation (r > 0.50, Appendix 1-Table 2) as some evidence from genomic studies suggests that variable importance measures may show a bias towards correlated predictor variables (Nicodemus et al., 2010). So, our final models included seasonality in temperature and precipitation as representatives of climate heterogeneity; velocity of climate change since the LGM and the MH to the present as representatives of past climate change; plate tectonics; mean elevation representing mountains; and human land use at the present, 5000 and 2000 years ago. Human land use 5000 and 2000 years ago showed a large correlation (r = 0.87), so we decided to perform separate models for each time period. Globally the correlation between human land use at present and 2000 years ago is relatively low (r = 0.29), however we also decided to perform independent models for both predictors because for certain bioregions both human land use at present and 2000 years ago can be highly correlated. We also ran additional models with the 536 extant species that have undergone some modification in their distribution range due to anthropogenic pressures. Finally, the algorithm used in the process of regionalization only uses species presence-absences, which means that humans can only have a role if present and if causing local extinctions, thereby changing the amount of shared species between grid cells. At the largest biogeographical scale two bioregions, Madagascar and Australia, are characterised by an extremely low historical human land use that could influence results. Therefore, we also ran the models excluding Australia and Madagascar. Random forest models were computed using the R package ‘*RandomForest*’ (Liaw and Wiener, 20002).

### Calculating global and local predictor importance

We measured variable importance using the mean decrease in accuracy, which is obtained by permuting randomly each variable and assessing the decrease in classification accuracy of the model (Liaw and Wiener, 2002). Thus, the more the accuracy of the random forest decreases due to the permutation (or exclusion) of a single variable, the more important that variable is considered, i.e., variables with a large mean decrease in accuracy are more important for classification of the data.

Random forest also calculates the local variable importance, which defines the importance of a variable in the classification of a single sample (grid cell here) and therefore shows a direct link between variables and samples (Touw et al., 2012). The local importance score is derived from all trees for which the sample was not used to train the tree (i.e. its value is OBB). The percentage of correct votes for the correct class in the permuted OBB data is subtracted from the percentage of votes for the correct class in the original OOB data to assign a local importance score for the variable for which the values were permuted. The score reflects the impact on correct classification of a given sample: negative, 0 (the variable is neutral) and positive. Given that local importance values are noisier than global importance ones we run the same classification 5 times (5 per biogeographical scale) and averaged the local importance scores to obtain a robust estimation of local importance values (Touw et al., 2012). Since one value per cell was obtained, these were used to create maps of the local importance of present and past human land use. However, note that the values of local importance do not distinguish between areas where the human land use was high and areas where it was negligible, i.e., both are equally valuable for the classification analysis. So, to help in the interpretation we also represent in maps the local importance value for those grid cells where the percentage of land used was greater than 10%.

All analyses were performed in the R Version 3.6.1 (R Core Team 2019).

## Data availability

All Data supporting this article will be deposited in Dryad upon acceptance.

## Acknowledgements

This work was supported by the project 2020/125-US/JUNTA/FEDER EU (Programa Operativo FEDER/Junta de Andalucía 2014-2020) to MR, and by CGL2017-83045-R AEI/FEDER EU (Agencia Estatal de Investigación from the Ministry of Economy, Industry and Competitiveness, Spain co-financed with FEDER) to ER.

