## Supplementary figures and tables for "Global biogeographical regions reveal a signal of past human impacts"

### Appendix 1

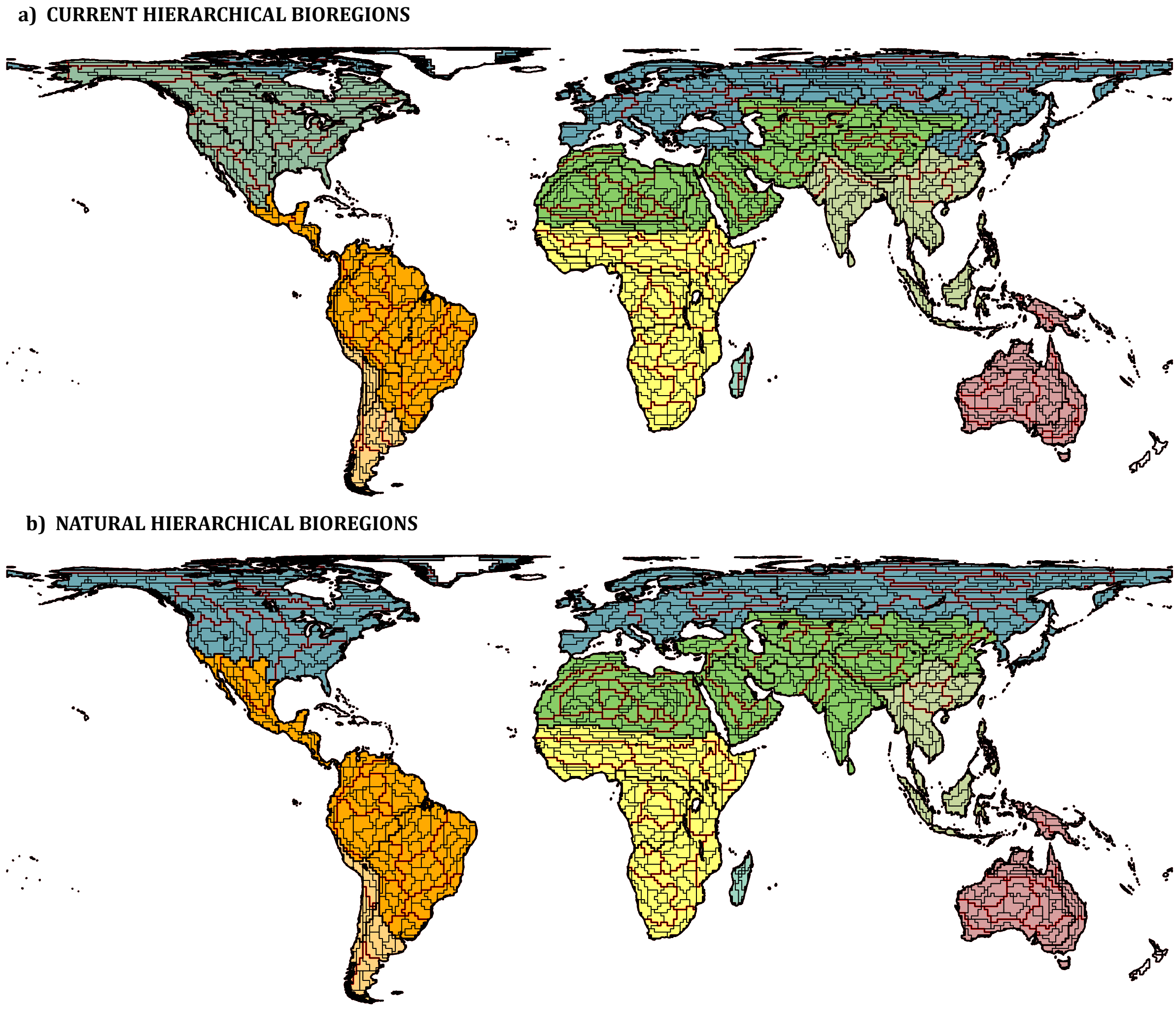


**Figure 1. Nested global bioregionalizations for non-volant terrestrial mammals.** The hierarchical application of the affinity propagation algorithm resulted in a nested global bioregionalization containing for the current hierarchical bioregions (a): 9 (filled colours), 27 (black boundaries), 141 (red boundaries) and 1128 (grey boundaries) bioregions from the largest extant realms to the smallest obtained bioregions; and for the natural hierarchical bioregions (b): 8 (filled colours), 25 (black boundaries), 121 (red boundaries), and 1051 (grey boundaries) bioregions.

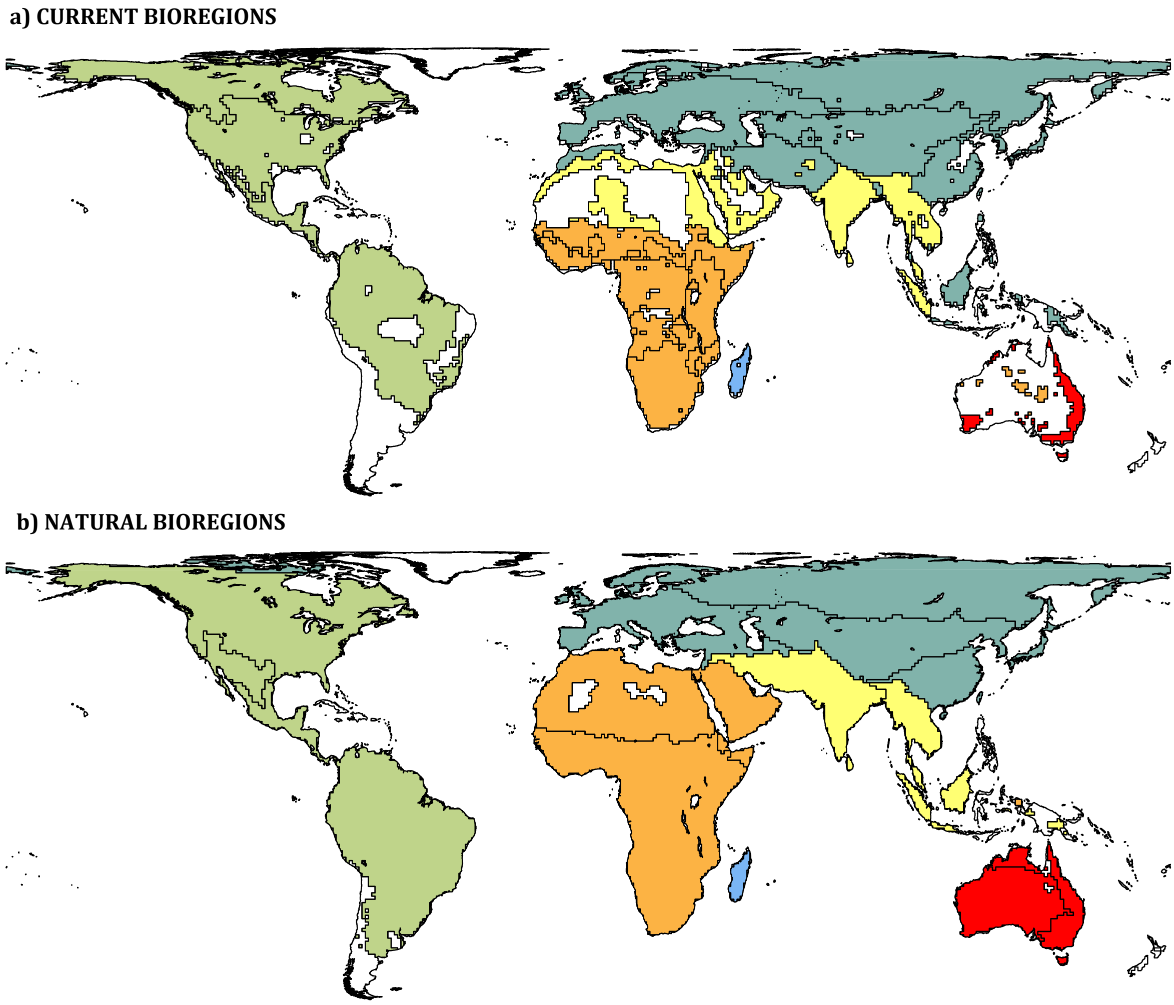


**Figure 2. Broader regions and medium-sized bioregions for the 536 extant non-volant terrestrial mammals whose distribution ranges have shifted due to human actions.** The affinity propagation clustering algorithm resulted in 6 broader bioregions (filled colours), and 18 subregions (black boundaries) for (a) Current bioregions; and 6 broader bioregions (filled colours) and 17 subregions (black boundaries) for (b) Natural bioregions. Although not represented in the map, the clustering algorithm resulted in 93 and 1333 smaller regions for the current, and 98 and 997 smaller regions for the natural bioregions.

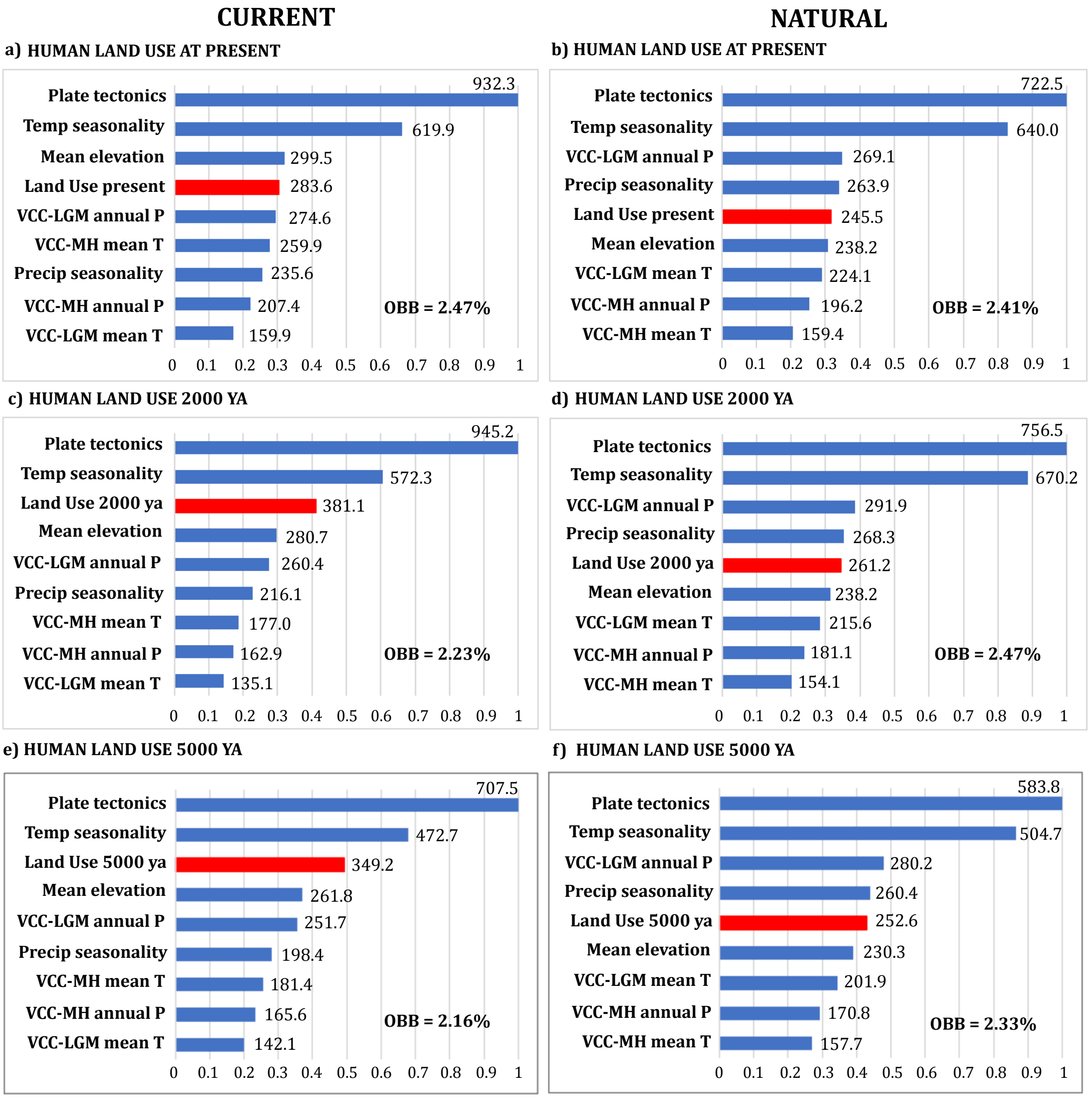


**Figure 3. Ranking of importance values for the drivers of taxonomic differentiation for the current and natural broader bioregions without Australia and Madagascar.** Above panels show importance values from models with human land use at present for (a) current and (b) natural bioregions, while the panels below show importance values from models with human land use 2000 years ago for (c) current and (d) natural bioregions, and human land use 5000 years ago for (e) current and (f) natural bioregions. Importance was measured by the drop-in classification accuracy after predictor randomization in random forests of 5000 trees. Higher values of mean decreased in accuracy indicate variables that are more important to the classification. OBB (Out-of-bag) represents the percentage of cells misclassified in each model. VCC = velocity of climate change; MH = Mid-Holocene; LGM = Last Glacial Maximum; annual P = annual precipitation; mean T = mean annual temperature; ya = years ago.

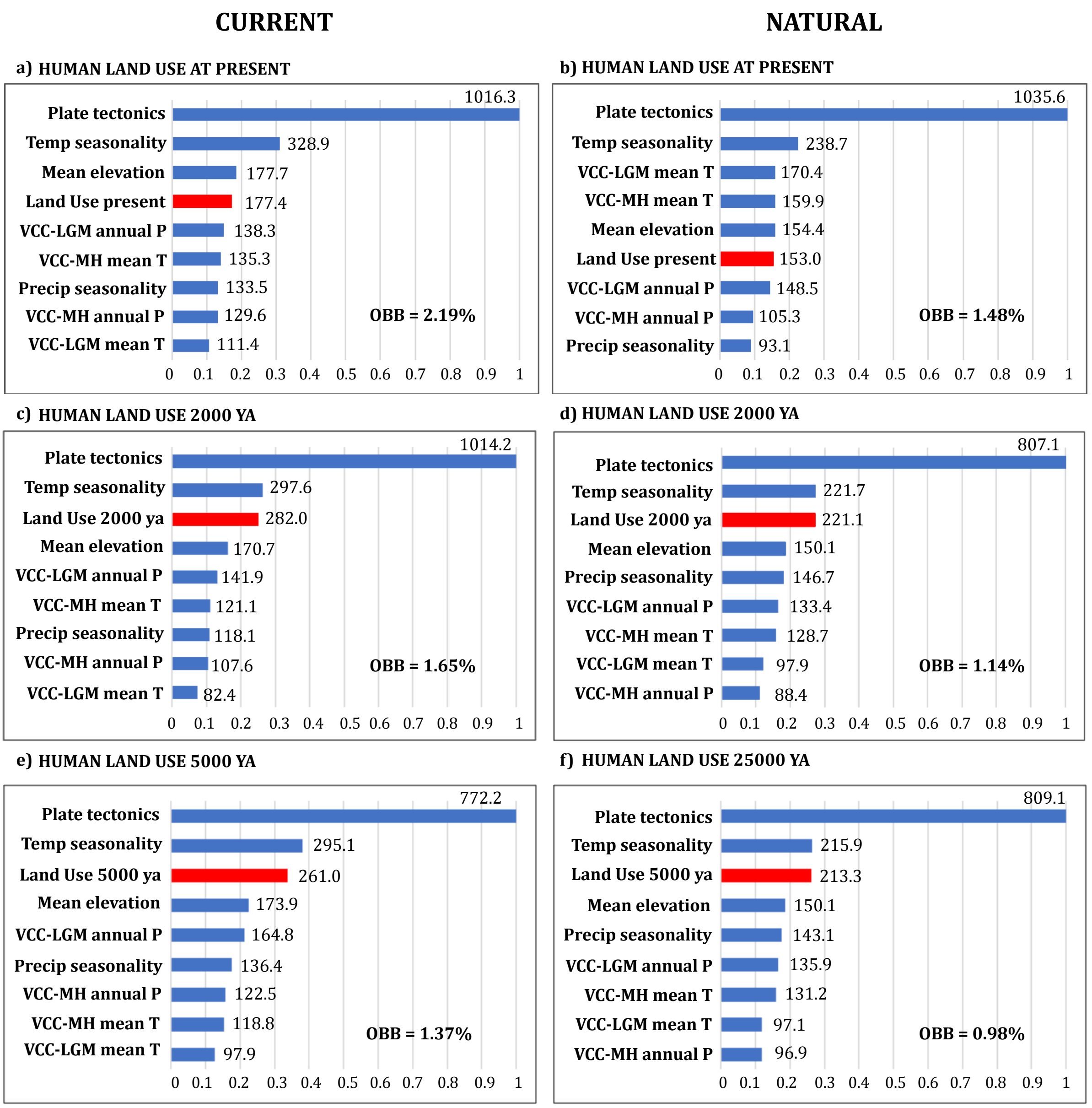


**Fig. S4. Ranking of importance values for the drivers of taxonomic differentiation for the current and natural broader bioregions for the 536 extant non-volant terrestrial mammals whose distribution ranges have shifted due to human actions.** Above panels show importance values from models with human land use at present for (a) current and (b) natural bioregions, while the panels below show importance values from models with human land use 2000 years ago for (c) current and (d) natural bioregions, and human land use 5000 years ago for (e) current and (f) natural bioregions. VCC = velocity of climate change; MH = Mid-Holocene; LGM = Last Glacial Maximum; annual P = annual precipitation; mean T = mean annual temperature; ya = years ago.

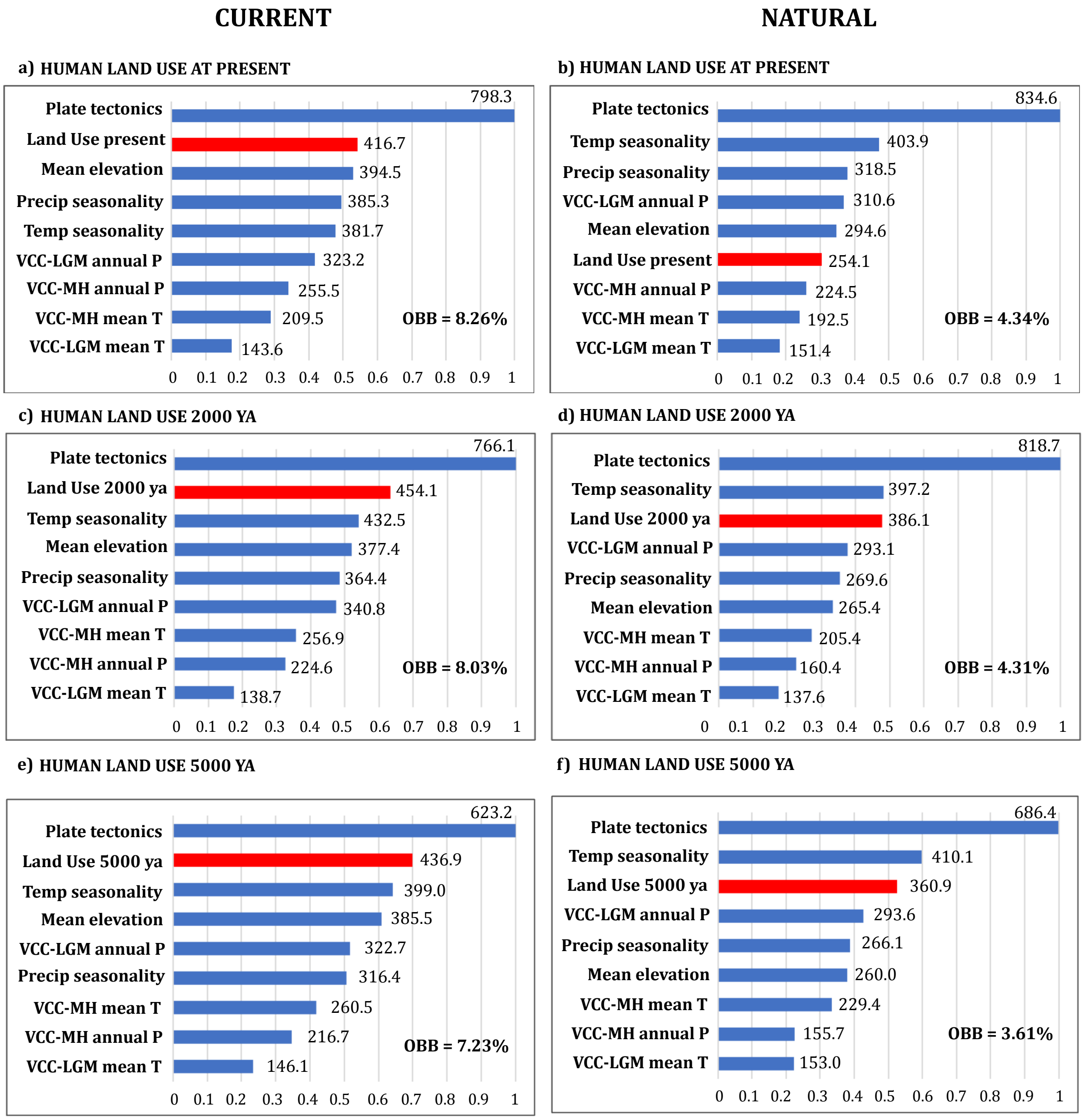


**Figure 5. Ranking of importance values for the drivers of taxonomic differentiation for the current and natural medium-sized subregions for the 536 extant non-volant terrestrial mammals whose distribution ranges have shifted due to human actions.** Above panels show importance values from models with human land use at present for (a) current and (b) natural bioregions, while the panels below show importance values from models with human land use 2000 years ago for (c) current and (d) natural bioregions, and human land use 5000 years ago for (e) current and (f) natural bioregions. VCC = velocity of climate change; MH = Mid-Holocene; LGM = Last Glacial Maximum; annual P = annual precipitation; mean T = mean annual temperature; ya = years ago.

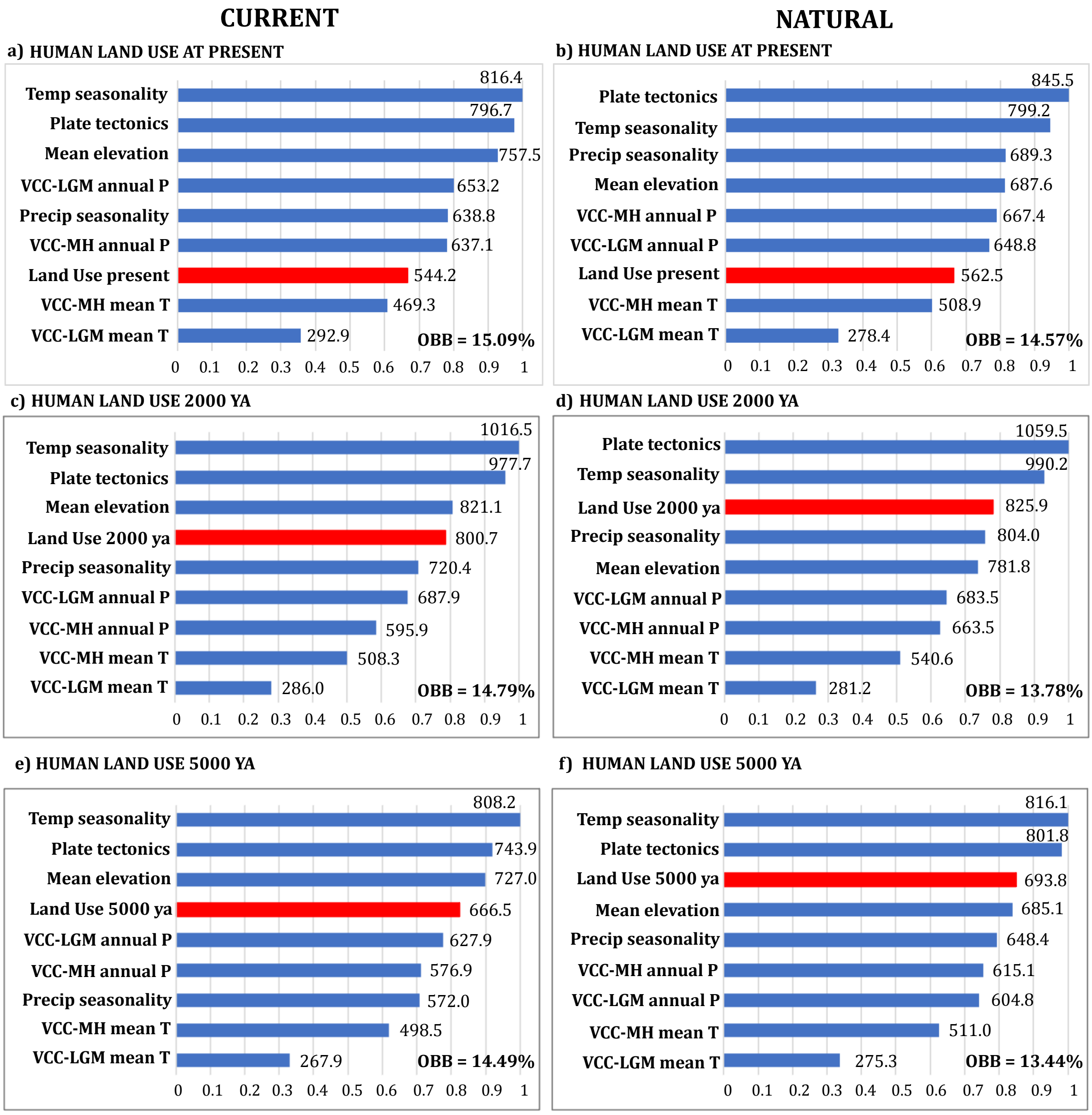


**Figure 6. Ranking of importance values for the drivers of taxonomic differentiation for the current and natural small subregions.** Above panels show importance values from models with human land use at present for (a) current and (b) natural bioregions, while the panels below show importance values from models with human land use 2000 years ago for (c) current and (d) natural bioregions, and human land use 5000 years ago for (e) current and (f) natural bioregions. VCC = velocity of climate change; MH = Mid-Holocene; LGM = Last Glacial Maximum; annual P = annual precipitation; mean T = mean annual temperature; ya = years ago.

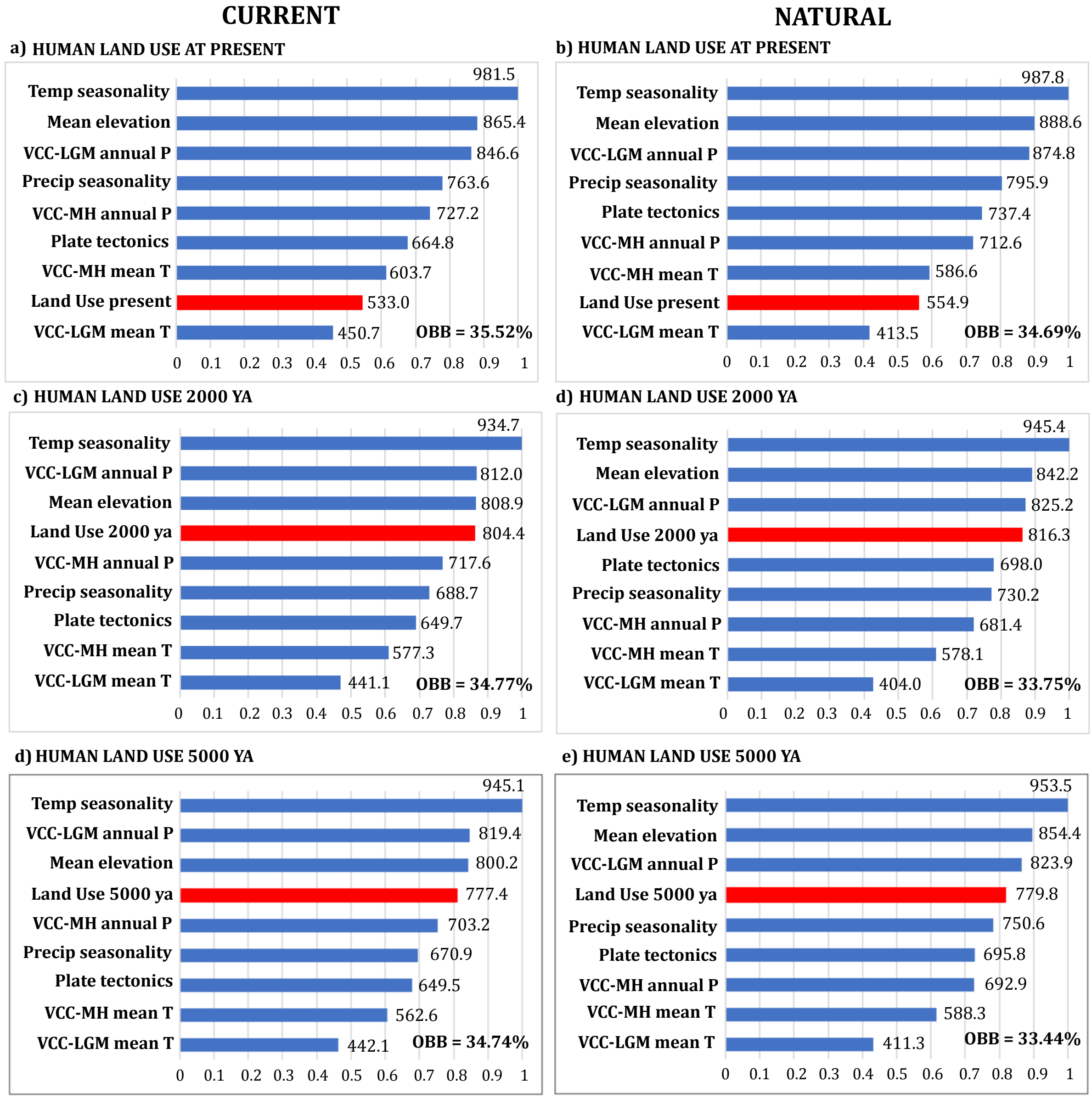


**Figure 7. Ranking of importance values for the drivers of taxonomic differentiation for the current and natural smaller subregions.** Above panels show importance values from models with human land use at present for (a) current and (b) natural bioregions, while the panels below show importance values from models with human land use 2000 years ago for (c) current and (d) natural bioregions, and human land use 5000 years ago for (e) current and (f) natural bioregions. VCC = velocity of climate change; MH = Mid-Holocene; LGM = Last Glacial Maximum; annual P = annual precipitation; mean T = mean annual temperature; ya = years ago.

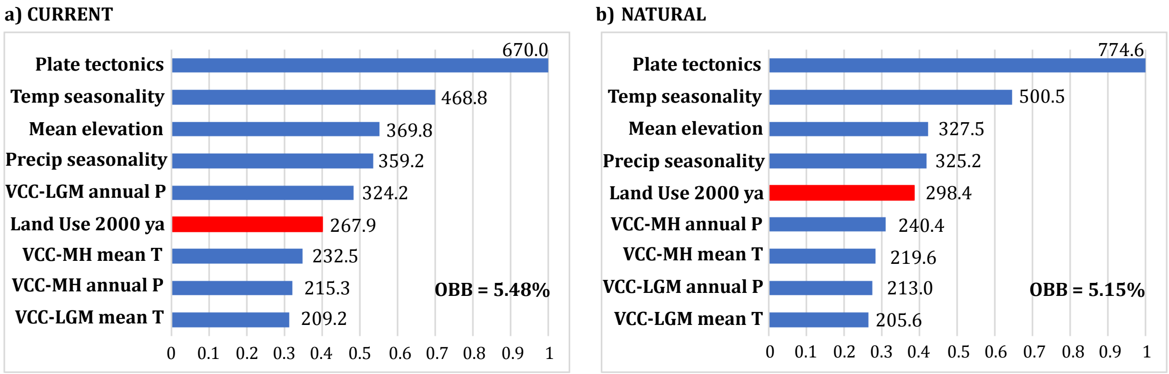


**Figure 8. Ranking of importance values for the drivers of taxonomic differentiation for the current (a) and natural (b) medium-sized subregions when subregions without human land use 2000 years ago are removed from the analysis.** Subregions removed were Madagascar, the two Australian subregions, the Sahara Desert (only for the current subregions), the Tibetan plateau, and the American Arctic. VCC = velocity of climate change; MH = Mid-Holocene; LGM = Last Glacial Maximum; annual P = annual precipitation; mean T = mean annual temperature; ya = years ago.

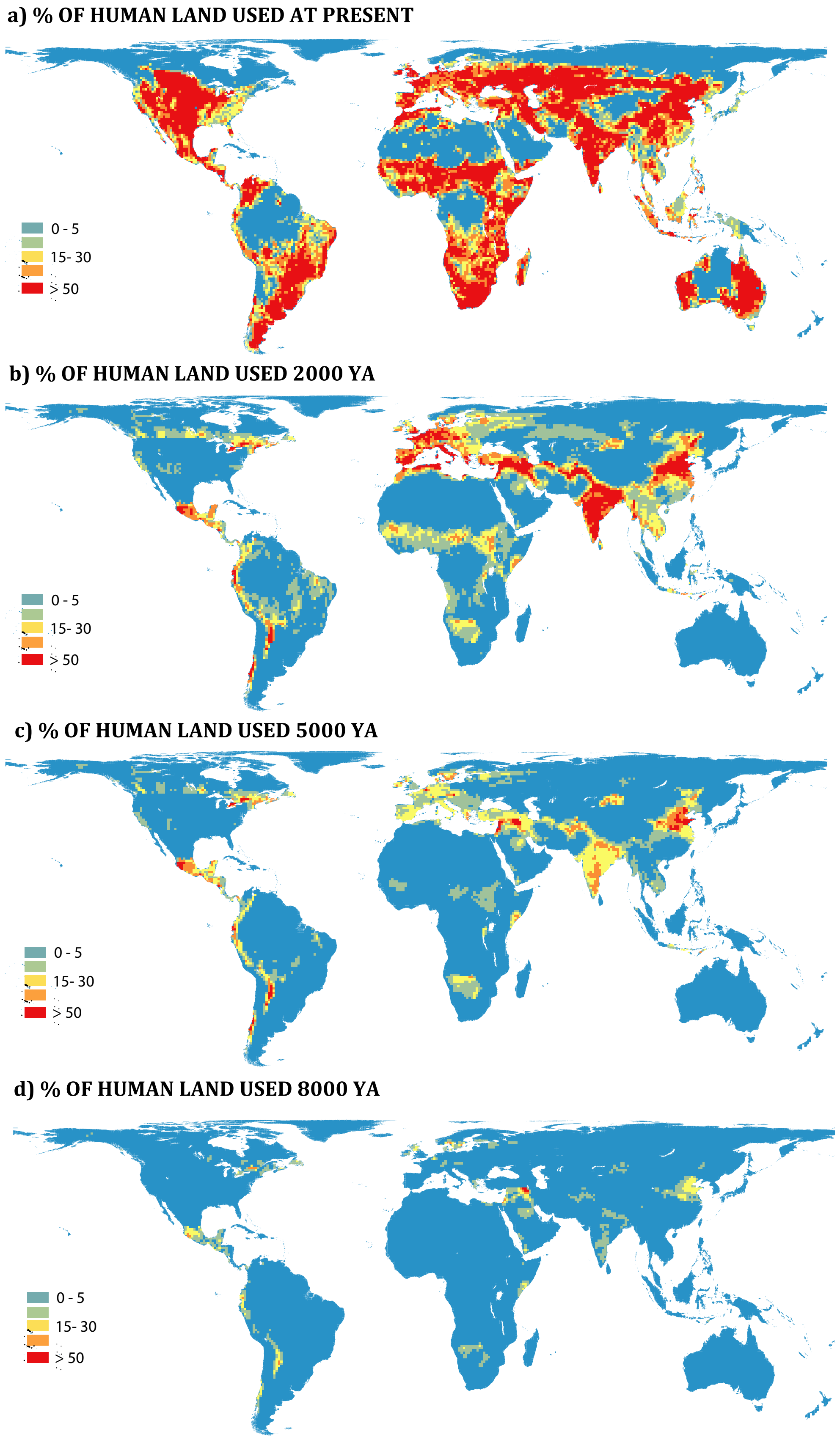


**Figure 9. Percentage of human land used at present, 2000, 5000 and 8,000 years ago.**

**Table 1. Jaccard index for current and natural hierarchical bioregions built with the 536 extant mammals whose distributions ranges have shifted due to human impact.** Jaccard index assesses the degree of similarity between clusters (bioregions) and ranges between 0 (no similarity) and 1 (perfect match). The number of bioregions obtained during each of the bioregionalization processes is also provided.

| CURRENT | Jaccard coefficient | NATURAL |
|---|---|---|
| 6 broader realms | 0.48 | 6 broader realms |
| 18 subregions | 0.33 | 17 subregions |
| 93 small bioregions | 0.20 | 98 small bioregions |
| 1333 smaller bioregions | 0.19 | 997 smaller bioregions |

**Table S2. Pearson's correlations between continuous determinants.** MAT = Mean annual temperature; VCC = velocity of climate change; MH = Mid-Holocene; LGM = Last Glacial Maximum; ya = years ago.

| | Elevation | MAT | Annual precip | Temp season | Precipit season | VCC-LGM Temp | VCC-MH Temp | VCC-LGM Precip | VCC-MH Precip | Land use present | Land use 2000 ya | Land use 5000 ya | Land use 8000 ya |
|---|---|---|---|---|---|---|---|---|---|---|---|---|---|
| Elevation | 1 | | | | | | | | | | | | |
| MAT | -0.21 | 1 | | | | | | | | | | | |
| Annual precipitation | -0.15 | 0.35 | 1 | | | | | | | | | | |
| Temp seasonality | -0.007 | -0.84 | -0.55 | 1 | | | | | | | | | |
| Precipit seasonality | 0.21 | 0.30 | -0.18 | -0.18 | 1 | | | | | | | | |
| VCC-LGM Temp | -0.27 | -0.41 | -0.12 | 0.44 | -0.29 | 1 | | | | | | | |
| VCC-MH Temp | -0.21 | 0.13 | 0.03 | -0.06 | 0.09 | 0.32 | 1 | | | | | | |
| VCC-LGM Precip | -0.17 | -0.16 | 0.33 | 0.04 | -0.19 | 0.37 | 0.04 | 1 | | | | | |
| VCC-MH Precip | -0.09 | 0.36 | 0.24 | -0.36 | 0.37 | -0.17 | 0.19 | 0.11 | 1 | | | | |
| Land use present | 0.07 | 0.20 | -0.06 | -0.10 | 0.13 | -0.08 | -0.05 | -0.12 | 0.13 | 1 | | | |
| Land use 2000 ya | -0.0007 | 0.12 | 0.07 | -0.15 | 0.13 | -0.01 | -0.01 | -0.08 | 0.04 | 0.29 | 1 | | |
| Land use 5000 ya | 0.016 | 0.07 | 0.05 | -0.06 | 0.11 | -0.04 | -0.03 | -0.04 | -0.01 | 0.22 | 0.87 | 1 | |
| Land use 8000 ya | 0.006 | 0.06 | 0.008 | -0.04 | 0.08 | -0.01 | -0.03 | -0.04 | -0.03 | 0.13 | 0.63 | 0.86 | 1 |
